# Topological and Structural Plasticity of the single Ig fold and the double Ig fold present in CD19

**DOI:** 10.1101/2021.06.04.447059

**Authors:** Philippe Youkharibache

**Affiliations:** Cancer Data Science Laboratory, National Cancer Institute, NIH, Bethesda MD 20814, USA

## Abstract

The Ig-fold has had a remarkable success in vertebrate evolution, with a presence in over 2% of human genes. The Ig-fold is not just the elementary structural domain of antibodies and TCRs, it is also at the heart of a staggering 30% of immunologic cell surface receptors, making it a major orchestrator of cell-cell-interactions. While BCRs, TCRs, and numerous Ig-based cell surface receptors form homo or heterodimers on the same cell surface (in cis), many of them interface as ligand-receptors (checkpoints) on interacting cells (in trans) through their Ig domains. New Ig-Ig interfaces are still being discovered between Ig-based cell surface receptors, even in well known families such as B7. What is largely ignored however is that the Ig-fold itself is pseudo-symmetric, a property that makes the Ig-domain a **versatile self-associative 3D structure** and may in part explain its success in evolution, especially through its ability to bind in cis or in trans in the context of cell surface receptor-ligand interactions. In this paper we review the Ig domains tertiary and quaternary pseudo symmetries, with a particular attention to the newly identified **double Ig fold** in the solved CD19 molecular structure to highlight the underlying fundamental folding elements of Ig domains, i.e. Ig protodomains. This pseudosymmetric property of Ig domains gives us a decoding frame of reference to understand the fold, relate all Ig-domain forms, single or double, and suggest new protein engineering avenues.

## INTRODUCTION

### Tertiary Pseudo symmetry of the Ig fold

We previously established that ca. 20% of known protein folds/domains are pseudo-symmetric (Myers-Turnbull et al. 2014) and that in each structural class (Lo Conte et al. 2000) the most diversified fold exhibits pseudo symmetry, suggesting a link between symmetry and evolution. In particular, two classes of folds show a higher proportion of pseudo-symmetric domains: membrane proteins, with for example GPCRs (Youkharibache, Tran, and Abrol 2020), and beta folds, with chief among them the Ig-fold (Youkharibache 2019). The Ig-fold is present in over 2% of human genes in the human genome (Lander et al. 2001) and it is overly represented in the surfaceome/immunome (Bausch-Fluck, Milani, and Wollscheid 2019; Díaz-Ramos, Engel, and Bastos 2011). Beyond antibodies, B-cell and T-cell receptors and coreceptors, the Ig-domain is present in a very large number of T cell costimulatory and coinhibitory checkpoints that regulate adaptive immunity with, in particular, the CD28 family of receptors containing the well known CTLA-4 and PD-1 receptors and their ligands from the B-7 family (Sharpe 2017; J.-C. D. Schwartz et al. 2002; Sharpe and Freeman 2002). Overall, the Ig-fold accounts for a staggering 30% of cell surface receptors’ extracellular domains (Díaz-Ramos, Engel, and Bastos 2011), making it a major orchestrator of cell-cell-interactions. What is especially remarkable with Ig-domains is their ability to interact, i.e. self associate, in both cis and trans trough cell surface receptor-receptor or receptor-ligand interactions. The very notion of cell surface receptor vs. ligand is arbitrary as Ig-domains are at the heart of a very elaborate network regulating immune response through Ig-Ig interactions in cis and in trans (Held and Mariuzza 2011; Chaudhri et al. 2018; Nishimura et al. 2021; Back et al. 2009; Zhao et al. 2019; Claus et al. 2019; Yang Liu et al. 2019; Q. Wang et al. 2019). A reason for self-interaction in cis or trans lies in its very structure: the Ig-fold is pseudosymmetric (Figure 1). While quaternary symmetry of Ig-domain based complexes is well known, the Ig tertiary structure pseudosymmetry is largely ignored, and we will review this property in both single Ig-domains and the recently solved CD19 structure with a novel double Ig-fold, a remarkable pseudosymmetrical protein architecture (Figure 4).

**Figure 1.**
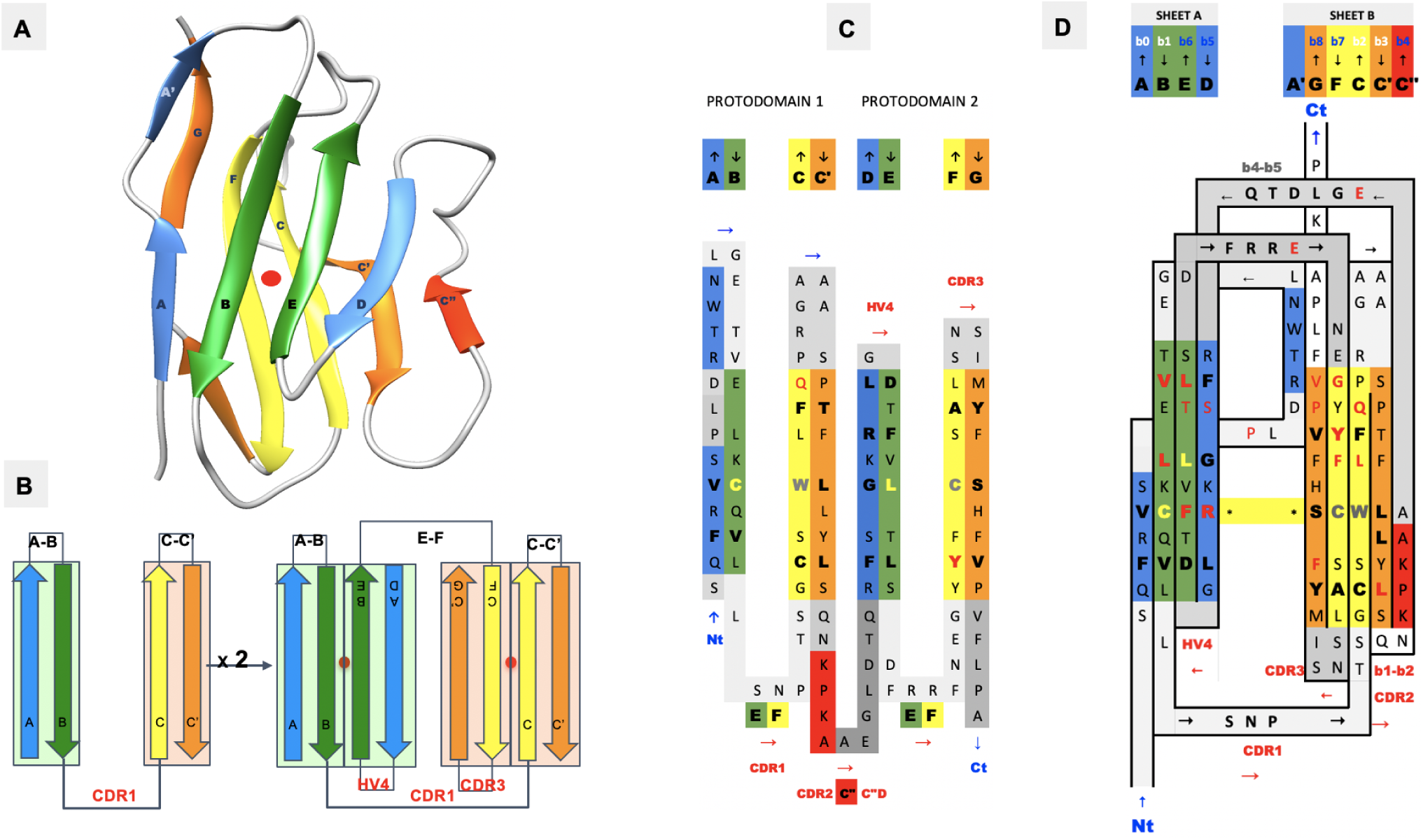
IgV domain deconstruction into pseudo-symmetric protodomains with an inverted topology. **A)** IgV domain - color scheme 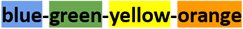 associated with each of the individual strands protodomain 1 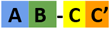 and protodomain 2 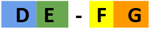 that align between 1 and 2A in most IgVs. and assemble pseudosymmetrically with a C2 axis of symmetry perpendicular to the paper plane. **B)** This corresponds to an inverted topology (using a membrane protein nomenclature) between the two protodomains. **C)** They invert through the linker **[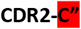 strand-C”D loop]. D)** The resulting IgV topology shows the self complementary assembly of the protodomains through their central strands 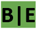 and 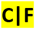 strands. Symmetry breaking occurs through the 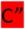 strand and through the 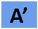. In IgVs, unlike IgCs, the 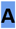 strand splits in two through a proline or glycines and participates to the two sheets 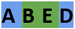 and 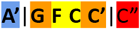. In figure A) we use PDBid 2ATP where the A’ strand is well formed. In D) the sequence/topology map for the CD8 sequence iCn3D link for CD8 (PDBid 1CD8).

#### Pseudo symmetry and ancient evolution of the Ig fold

A pseudo symmetric domain is formed of two or more protodomains, according to an accepted duplication-fusion mechanism (Lee and Blaber 2011), and multiple examples of highly diversified structural folds have been known for a long time (Youkharibache 2019). Structurally, it is important to realize that the knowledge of protodomains and symmetry operators define a pseudo-symmetric protein domain entirely, apart from a **variable linker region**, most often short, chaining protodomains within a domain (Youkharibache 2019). Interestingly, a pseudo-symmetric domain is a tertiary structure that can also be considered a pseudo-quaternary structure, and can be analyzed according to symmetry groups (C2 or higher C3, C4 … D2, …Dn), with a C2 symmetry prevalence in the known structural protein universe ([PDB symmetry statistics](https://www.rcsb.org/search/browse/symmetry)). The symmetric assembly of (chained) protodomains is a general property found in biopolymers beyond proteins, as in RNA riboswitches for example (Youkharibache 2019; Jones and Ferré-D’Amaré 2015), demonstrating a general duplication mechanism in biopolymers with symmetric self assembly of the duplicated parts, beyond the realm of proteins, and already present in the RNA world when considering the ribosomal peptidyl transferase center (PTC) (Bashan et al. 2003) as a remnant of an ancient proto-ribosome.

Remarkably, thinking of an ancient peptide/protein world, a very early investigation of Immunoglobulins in sequence space (Urbain 1969) suggested that “Each part of the immunoglobulin chains (Fd, Fc, v, c) derives from the repetition of a smaller ancestral sequence (of approximately 20 amino acids) [...] The following scheme of evolution is suggested: In the first steps, we shall speak only of peptides and not of genes, since it is not known if ancestral peptides were coded by nucleotide sequences. The process started with a primordial peptide which, by a special mechanism, doubled its length and became (pseudo)symmetrical”. One can only be impressed by such an early and insightful analysis, since we are now able to discern pseudosymmetry in Ig domains structurally, and delineate protodomains. Further efforts tried to systematize sequence based pseudo symmetrical analysis of Ig domains (Yanzhao Huang and Xiao 2007), but when comparing to 3D structure based conservation of protodomains, sequence alignment matches do not exactly coincide with structure alignment matches. This is typical of sequence-structure internal symmetry detection (Youkharibache 2019; Youkharibache, Tran, and Abrol 2020; Myers-Turnbull et al. 2014), and if sequence matches are real, they imply a shift in relative position of sequence in structure, the evidence of which is difficult to demonstrate, an interesting observation nonetheless, albeit anecdotal.

Another early study suggested some “evidence for an ancestral immunoglobulin gene coding for half a domain” (Bourgois 1975) based on sequence considerations, postulating the half domains would correspond to strands AB-CC’ and DE-FG and assemble as an IgV domain, using an accepted nomenclature of Ig domain strands (Lesk and Chothia 1982; Williams and Barclay 1988). Later this delineation was supported on structural considerations by McLachlan (McLACHLAN 1980). These elements are not well known, but have been reviewed in the literature (Williams and Barclay 1988) and are supported by our observations based on structural pseudo symmetry of single Ig domains (Youkharibache 2019). They are also supported by an observed swap of the GFCC’ protodomain between IgV domains in a dimeric form of CD2 (Murray et al. 1995, 1998) and, as we shall see in the following in the case of CD19, the AB-CC’ and DE-FG protodomains interdigitate to form a novel double Ig-fold (Teplyakov et al. 2018).

##### Converging vs. diverging evolution

For many Ig domains, and even more when also considering structurally similar domains such as FN3 or Cadherins, the sempiternal question comes back: are these domains evolutionarily related? Do these domains have a common ancestor? As for many highly diversified superfolds domains (Orengo and Thornton 2005; Myers-Turnbull et al. 2014; Youkharibache 2019; Youkharibache, Tran, and Abrol 2020) the question is relevant. It is especially relevant as it relates to the various topological forms of Ig-domains. The protodomain (half-domain) hypothesis tends to support the IgV domain as the ancestral form in a divergent evolution scenario. Arguments on the origin of the Ig domain have been partially reviewed in the literature (Williams and Barclay 1988; Oreste, Ametrano, and Coscia 2021). In the following we analyze the structural and topological domains in light of the Ig-fold pseudosymmetry and a possible parallel evolution of these topological domains.

### Quaternary pseudo symmetry of Ig-domains assemblies: Ig-domains dimerization

Let’s briefly review the pseudo symmetric assembly of Ig-domains. Twofold (C2) quaternary symmetry has been demonstrated in the very first structural studies on Ig-domains dimers, especially of IgV-domains (Epp et al. 1974, 1975; Poljak et al. 1974). The IgV-IgV dimerization interface using the **GFCC’** sheet was observed in the first Fab structure solved (7FAB) (Saul, Amzel, and Poljak 1978) and can be considered canonical. It is also found in homodimers such as CD8aa (1CD8) (Leahy, Axel, and Hendrickson 1992), and was in fact already observed in the Bence Jones protein, the very first structure deposited in the PDB (1REI) (Epp et al. 1975), forming a VL-VL interface through that same **GFCC’** sheet. In antibodies and antibody fragments such as Fabs or scFvs, pseudosymmetric interfaces are formed by VH-VL domain pairing. Constant domains CH-CL, on the other hand use the opposite sheet **ABED** of the Ig domain to dimerize. In some cases, VH domains can also pair through that ABED sheet, as in Fab-dimerized glycan-reactive antibodies (Acharya et al. 2020).

Multiple different dimeric interfaces have been characterized in particular between IgV domains at the N terminus of cell surface receptors and ligands, where a majority of them interact through their GFCC’ sheets but in a wide variety of orientations. Other pseudo symmetric parralel dimer interfaces for CTLA-4 or CD28 homodimers on T-cell surfaces for example use instead their A’G strands and their Hinges beyond their G strand to dimerize, and in doing so free the GFCC’ sheet of both IgV domains to interact with other IgV domains in either cis or trans; in particular in trans using using the GFCC’ sheet of their ligands CD80 or CD86, that themselves homodimerize in cis using their strands C”D [see Figure 2], and/or in cis as in the case of CD80/PD-1. It is beyond the scope of this paper to review all possible interfaces, but let us note nonetheless that the canonical GFCC’ interface can be considered, in most cases, a “cis” interface, as the domains are parralel, positioning both domains C terminus towards a given cell membrane, while “trans” IgV-IgV interfaces also using the GFCC’ sheet can be formed from opposite cell membranes in antiparralel, for example between the CTLA-4 (or CD28) receptor on the surface of T-cells with their natural ligands CD80 or CD86 (J. C. Schwartz et al. 2001) on the surface of targeted cells [see Figure 2]. This is not a rule however, due to the flexibility of some surface receptors’ hinges as in the case of PD-1|PD-L1 or PD1|PD-L2 interactions, where IgV domains form a parallel interface while on opposite cell surfaces, resembling the canonical interface and implying an inversion of the PD-1 domain in binding a PD-L domains [(Freeman 2008; Lázár-Molnár et al. 2008; Zak et al. 2015). In all IgV interfaces, a quaternary C2 axis of pseudo symmetry is observed, but that axis varies with the relative orientation of IgV dimers. The characterization of the full set of interfaces formed by Ig-domains still awaits a review, but let us note that new Ig-Ig binding interfaces are still being discovered even in the best studied within the Ig superfamily such as the B7-family with CD86 or CD80 homodimers in cis, PD-L1/CD80 heterodimers in cis, using the same interface, CTLA-4/CD80 in trans PD-L1/PD-1 in trans using once again the same interface (Garrett-Thomson et al. 2020).

**Figure 2.**
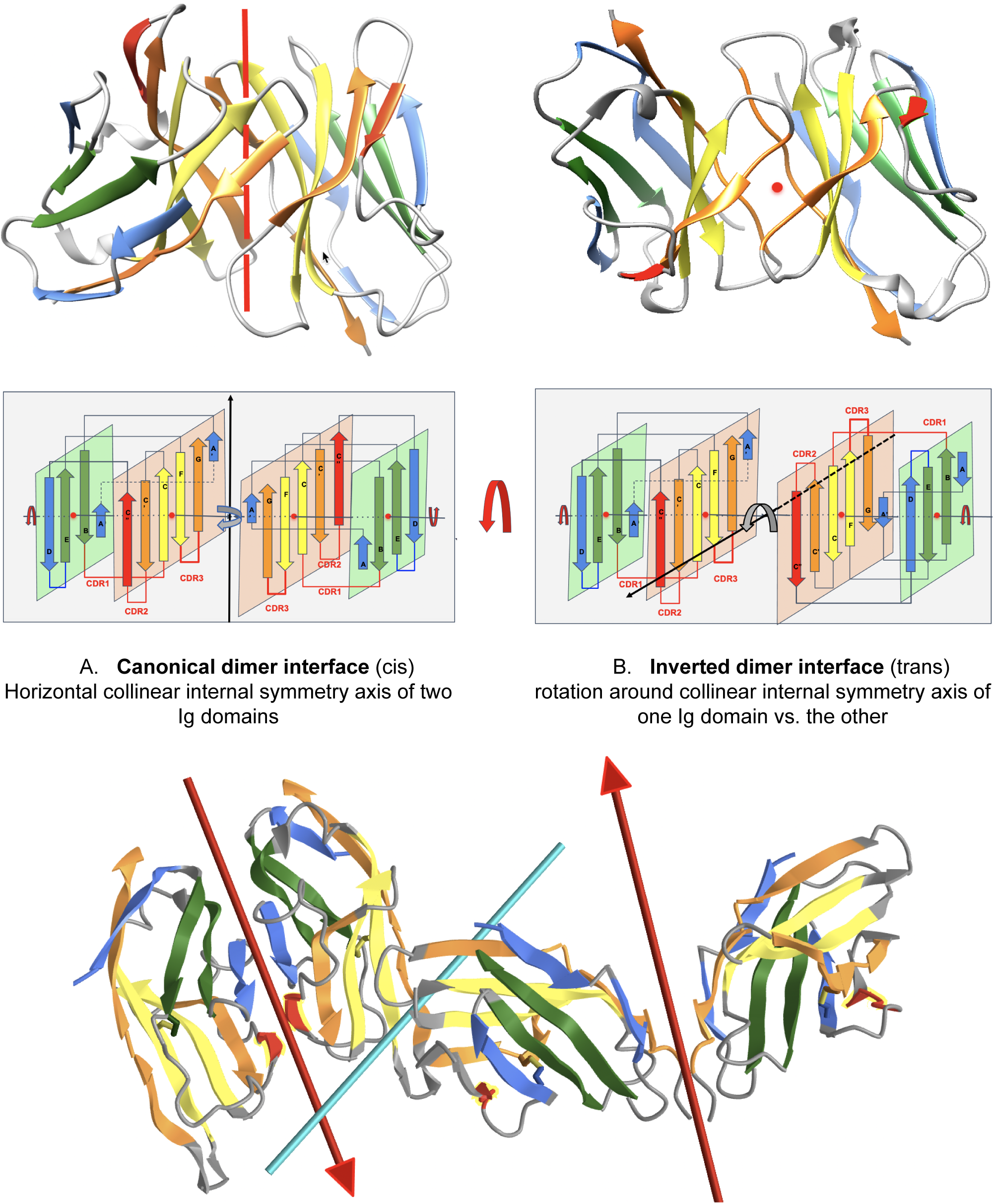
Parallel and antiparallel Ig domain dimerization interfaces. **A) Canonical dimer interface using the GFCC’ sheet:** The observed quaternary axis of symmetry is vertical. This is the classical interface of antibody VH-VL domains with the CDRs on the same side. It is also found in the Bence Jones protein as a VL-VL or CD8aa
as a homodimer, or CD8ab as a heterodimer. The Figure shows the structure of CD8aa interacting through the GFCC’ sheet in parralel. Its Topology/Sequence map can be seen in Figure 1 [iCn3D-CD8 parallel homodimer(1CD8)](https://structure.ncbi.nlm.nih.gov/icn3d/share.html?JcP3sd1gGfqXEBEM8). **B) Inverted dimer interface using the GFCC’ sheet:** The observed quaternary axis of symmetry is horizontal coming out of the page plane. It corresponds to a rotation around the collinear internal symmetry axis (horizontal left to right) of one Ig domain vs. the other. In this case CDRs are on opposite sides, the C terminus G strands are pointed in opposite directions. The Figure shows the structure of a VL-VL dimer interacting through the GFCC’ sheet in antiparallel iCn3D inverted homodimer (7JO8). As can be observed in the top drawings and in the 3D models [Links], the domains are tilted relative to each other (Lesk and Chothia 1982), this is not represented in the idealized models in the lower drawings. If one considers the tertiary C2 pseudo symmetry of individual domains, when they collinearize, the axes of symmetry cross the quaternary symmetry axis at the center of symmetry and form a higher pseudo symmetry group D2. In most complexes however the tertiary and quaternary axes do not cross exactly at the center, as in PD-1/PDL-1. This is an idealized model naturally, yet in homodimers, while we observe a quasi D2 symmetry however. The coloring 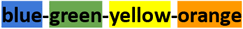 associated with each of the individual strands of protodomains 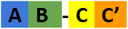 and 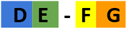 self assemble as an Ig-domain through their central green strands 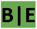 and yellow strands 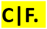 Symmetry breaking occurs on domain edges through the 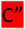 strand in red and through the 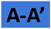 splitted strand. The 3D links for the Bence Jones protein with a canonical interface [iCn3D-1REI (VL-VL)](https://structure.ncbi.nlm.nih.gov/icn3d/share.html?RK2mWw4w3BfULb2H6) and [iCn3D-7JO8 inverted (VL-VL)](https://structure.ncbi.nlm.nih.gov/icn3d/share.html?eGmk1SeSjLyok3p37) and [comparison of the two with their respective symmetry axes and dimer interactions](https://structure.ncbi.nlm.nih.gov/icn3d/share.html?EWFzw5ZznZ8ZRsRV9). **C) CD86 dimer and CTLA-4 dimer interacting on opposite cell membrane surfaces using a variety of interfaces:** CD86 homodimerizes in cis using its strands C”D; CTLA-4 dimerizes in cis (similarly to CD28) on the opposite T-cell surface using strands A’ and G and most importantly the Hinge connecting the G strand to the transmembrane domain (forming a dimeric cystine bridge not shown in Figure but present in another 3D structure iCn3D - 3OSK), leaving in both CD86 and CTLA-4 their GFCC’ sheets to interact in trans iCn3D - 1I85 (CTLA-4/CD86).

To help with the understanding of Ig-Ig interfaces, some canonical homodimers such as CD8aa or VL-VL Bence Jones proteins, the tertiary C2 axis of symmetry combines with the quaternary C2 to lead to a quasi D2 symmetry group. This holds true even for VHVL or CD8ab heterodimers. Interestingly, some inverted homodimer interfaces are observed, where IgV domains are flipped (180 degrees) w.r.t. each other, resulting in a quaternary axis of symmetry rotated by 90 degrees, still orthogonal however to the tertiary C2 symmetry axis and an overall quasi D2 symmetry in considering the quaternary structure (see Figure 2). These structures are closer to idealized dimers and can help us understand a possible origin of the ubiquitous cis vs trans pairing of IgV domains, with parallel vs. antiparallel dimer interfaces. Using a membrane protein nomenclature, the latter is equivalent to an inverted (albeit quaternary) topology as opposed to a parallel one for canonical interfaces. In a canonical VH-VL antibody interface the CDR loops are on the same side, while in an inverted topology they would be on opposite sides. In Figure 2, we show an idealized representation comparing both dimer orientations and pseudo symmetries. Even in homodimers, IgV domains can not only adopt a parallel or inverted (flipped) orientation, they can adopt tilted orientations in between (D. B. Huang et al. 1994), all related to each other through a rotation around the tertiary C2 axis of symmetry. Further divergence from an idealized form of D2 symmetry, especially in receptor-ligand interactions, shifts these tertiary C2 symmetry axes away from collinearity. It is tempting to postulate that the tertiary pseudo symmetry of the Ig-domain may be the structural basis of its self association in either cis or trans, and for the latter in an inverted (quaternary) topology, yet not excluding a parallel one, as noticed before with PD-1/PD-L1. This is visualized in Figure 2 and beyond through the coloring scheme we have chosen to highlight symmetry with protodomain equivalence. (3D pictures are either from Chimera (Pettersen et al. 2004) or iCn3D (J. Wang et al. 2020). All 3D interactive graphics hyperlinks in the main text of figure legends use iCn3D and can be used directly by readers not only for visualization but also for further analysis).

In the following, we focus on the tertiary symmetries across all topological variants of Ig domains and dissect the formation of a new tertiary fold: the double Ig domain observed in CD19, which confirms the protodomain hypothesis.

## RESULTS

### Protodomain Evidence in single Ig-domains

As mentioned in the introduction, Nature may have used a protodomain duplication-fusion mechanism in a distant evolutionary past to build current day pseudo-symmetric domains, and we can gain insight in domain creation from an analysis of tertiary pseudo-symmetry of structurally characterized domains to look at domain creation and evolution in terms of their constituting parts. The Immunoglobulin fold is at the heart of a very large number of cell surface proteins of the immune system, beyond immunoglobulins themselves, and we have seen that the Ig-fold exhibits tertiary symmetry as well as quaternary symmetry, as in CD8 or VL-VL homodimer or the very well known antibody variable domains association: VH-VL.

#### The single Ig-fold pseudosymmetry and protodomains hypothesis

In Figure 3, we show schematically the topology of Ig domains in the Variable form (IgV), the Shark Variable form (VNAR) and the C1 Constant domain (IgC) [see more topological variants in Figure 6]. An Ig domain can be considered formed as a covalently linked dimer of two protodomains AB-CC’ and DE-FG using the standard strand terminology of Immunoglobulin domains (Lesk and Chothia 1982; Williams and Barclay 1988). Protodomains forming an Ig-domain usually align within 1 to 2 Angstrom RMSD range [Figure 6]. The two protodomains AB-CC’ and DE-FG are composed of two hairpin AB and CC’ for the first, DE and FG for the second, linked through BC and EF loops respectively, and whose relative orientation can be compared to an “inverted topology” (using a membrane protein nomenclature). We will refer to them with these denominations throughout the paper. Naturally Ig domains show more complexity with additional strands along these core central strands to form, in the IgV for example ABED and A’|GFCCC”’ sheets with the additional strands as we shall see. In figures, we use a protodomain spectrum coloring scheme: Blue-Green-Yellow-Orange for protodomain strands ABCC’ or DEFG and red for C” (see Figure 1).

**Figure 3.**
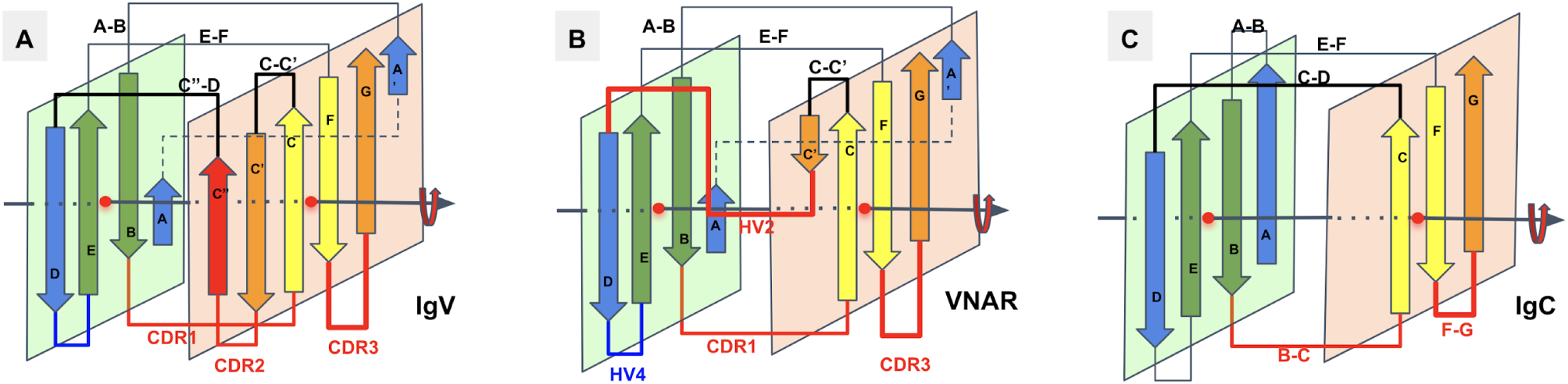
Ig domains Topologies for IgV, Shark VNAR, and IgC. **A)** In IgV domains the A strand, has a flexible hinge in the middle, usually a cis-proline or a stretch of glycines, and swaps the upper part of the strand (noted A’) to Sheet B. **B) VNAR** shows that same domain level organization with two protodomains, yet a much smaller inter protodomain linker, eliminating the linker’s secondary structure as present in IgV domains: the C” strand and the CDR2 loop. Instead a short HV2 is observed. In the literature, the C’ strand is usually included in the HV2 region, as it is extremely short. In addition a hydrophilic set of residues facing out on Sheet B (GF|CC’) rather than hydrophobic in IgV do not permit the formation of a symmetric dimer. Dimerization in crystal structures can be observed through HV2, yet they may not be relevant biologically. **C) IgC.** Here we consider only the IG C1-set, i.e. the antibody constant domain-like to exemplify an additional protodomain connectivity. In this case, the final domain is formed by a full 4-stranded AB|ED (Sheet A) vs a 3 stranded GFC Sheet B, and no C’ strand. Interestingly this enables the domain level dimerization through that 4-stranded Sheet A to form an 8-stranded barrel, as opposed to the IgV that uses the opposite Sheet B to form a 8-stranded dimer barrel.

One can reconcile all Ig forms (IgV, IgC, …) by considering an Ig domain formation through a pseudo symmetrical combination of protodomains with a variety of linking sequences that can form or remove strands on one sheet or the other. This opens the door to the interpretation of the formation of the diverse Ig topologies as a **parallel evolution process** leading to Ig-domains with variable topologies. A divergent evolution scheme would involve one domain formation with insertion or deletion of sequence elements between strands C and E leading to all topologies involving strands C’,C”, and D. This is similar to what we have described previously in pseudo symmetric polytopic membrane protein formation (Youkharibache, Tran, and Abrol 2020).

### Protodomain Evidence in the CD19 double Ig-fold

#### Characterization of the CD19 Ig domains in sequence and structural databases

Recently the structure of CD19 has been solved (Teplyakov et al. 2018). Prior to that knowledge, any textbook and sequence database would have presented CD19 as formed of two Ig domains of C2 topology in tandem, and in fact, this is still the case today in databases, where UNIPROT (P15391) and databases “confidently predict” CD19 extracellular region as composed of two Ig domains belonging to the IgC2-set (see discussion hereafter). From a sequence perspective they are not that far off, but the devil is in the details and sequence based Ig-domains are just not folding as two single Ig domains in tandem, but as one double Ig domain. Could a sequence analysis reveal a double Ig domain? Probably not, sequence matching algorithms clearly detect two Ig domains, however a detailed analysis shows that there are two areas unique to the CD19 extracellular domain sequence, first the two “Ig domains” are not connected in tandem by separated by a long linker, that we now know is structured as a small (inverter) domain (see Figure 4), and also a short “protodomain linker” within each of the “detected Ig-domains”. This is difficult to interpret for sequence matching algorithms using defined topological patterns (I-set, V-set, C1-set and C2-set). What is more surprising however is that an automated structural pattern matching algorithm (ECOD) would assign the double Ig domain as a V-Set followed by an I-set (Schaeffer et al. 2017).

**Figure 4.**
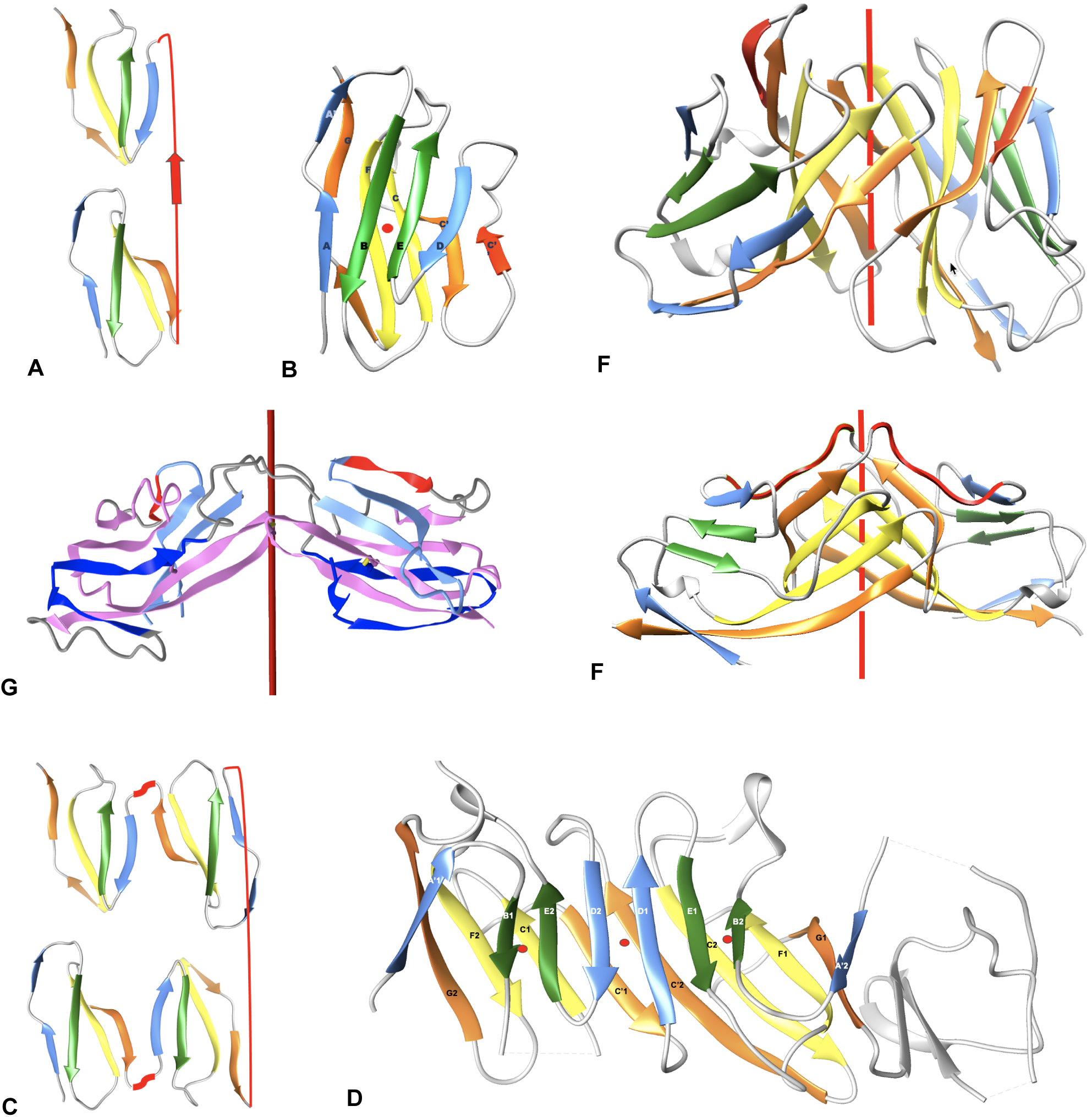
Single IgV and double IgV domain deconstruction. IgV dimers vs. double Ig domain (CD19) **A) Single IgV domain** schematic structural deconstruction **of a B) IgV domain**(iCn3D-1CD8): Two protodomains **AB-CC’** and **DE-FG** are fused in one domain in antiparallel, i.e. an “inverted-topology” in membrane protein terminology. See Figure 1 for details. **C) Double Ig domain CD19** schematic structural deconstructio**n** (in a CD19 domain the A chain is not present, only A’) **D) CD19 double Ig domain** (iCn3D-6AL5): the first two protodomains {AB-CC’ - DE-FG}**_1_** are chained together in a “parallel-topology” (with a short linker C’-D in red). The second protodomain pair {AB-CC’ - DE-FG}**_2_** assembles with the first pair pseudo-symmetrically through an “inverted topology”, thanks to a small intercalated domain linker (in gray). Local pseudo-symmetries between protodomains 1+4 and 2+3 are conserved, as for a regular IgV. The central axis of symmetry of the double Ig and the two local axes of symmetry of the composite Ig domains are shown as red dots perpendicular to the plane of the paper. Color shows local self-associations, at the strand level: as in regular or swapped Igs: through their green strands 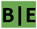 and and yellow strands 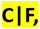 and in addition the double Ig assembles through the central strands 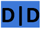 in blue and 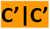 in orange, the 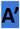 strands in both Igs are associated with the GFCC’ sheet (no A strand). **E) An IgV (canonical) CD8 dimer** (iCn3D-1CD8). The IgV dimer forms an 8-stranded central barrel two GF|CC’ sheets as homophilic CD8aa or heterophilic CD8ab dimers. Many Ig-based surface receptors use that interface, and so do antibodies in pairing VH-VL domains. **F) A swapped CD2 IgV-dimer** (iCn3D-1CDC) is observed between two IgV domains of CD2 that swap their second respective protodomains DE-FG to lead to a dimer where the first domain is composed of protodomains 1 (**AB-CC’**) and 4 (**DE-FG**) and the second Ig domain is composed of protodomain 3 (**DE-FG**) and 2 (**AB-CC’**). The linkers (in red) between protodomains 1-2 and 3-4 extend to bridge the two swapped IgV domains. C2 symmetry is preserved. **G) A swapped “tertiary dimer” (MXRA8)** (iCn3D-6JO8) - swapping the AB substructure resulting also in a pseudo symmetric tertiary structure in a head to head, resembling a quaternary swapped dimer as in F) but without forming a central barrel; see text for details. Coloring AB dark blue CC’ light blue, C” red, protodomain DE-FG magenta.

#### Protodomain Evidence in the CD19 double Ig-fold

The double Ig fold adopted by CD19 (Teplyakov et al. 2018) can be understood in terms of protodomains. In regular IgVs AB-CC’ (p1) and DE-FG (p2) protodomains are assembled in antiparallel with an internal C2 pseudosymmetry, through the characteristic {CDR2 loop + C’’ strand + C’’D Loop} structured “linker” [see Figure 4]. Shorter topologies (I-set, C1-set and C2-set) differ in the linker between protodomains (see Discussion below). It is easy to misidentify the strands C’ or D in the case of CD19 and to call for IgC2 domains. In CD19, strands C’ and D are observed and participate in consecutive protodomains AB-CC’ (p1) and DE-FG (p2) that are separated by a very short linker that forces them to remain parallel, unlike any known Ig-domain topology. This in itself implies a structural resilience of protodomains as they can associate in parallel, or invert. To reconcile these elements with the sequence analysis “the first Ig domain detected’’ adopts a parallel topology, and so does the second, something novel. Both “Ig-domains of parallel protodomain topology” as identified by sequence analysis can then assemble through a long linker that allows them to assemble/intertwine in an inverted topology (antiparallel with a C2 symmetry) juxtaposing their two BED sheets together and the two A|GFCC’ together as two long beta sheets facing each other [see Figure 4]

CD19 offers a higher complexity buildup than single Ig-domains in combining 4 Ig-protodomains to form an interdigitated pseudo symmetric double Ig-domain, a fold never observed before (Figure 4D). Crystallographers describe the structure they solved as a domain swap (Teplyakov et al. 2018), involving protodomains. CD19 topological innovation however results from a much more subtle and interesting folding. [A case of protodomain swap in CD2 is known](https://structure.ncbi.nlm.nih.gov/icn3d/share.html?AdsvRptE9BHqWnN56) and it is not the same as a CD19 fold [Figure 4F]. Noticeably, the authors at the time had hypothesized a possible protodomain duplication in the early evolution of Ig-domains (Murray et al. 1995), i.e. the half domain hypothesis described earlier (Bourgois 1975; McLACHLAN 1980), and even engineered higher order oligomers swapping protodomains (Murray et al. 1998). What Nature has achieved however in the case of CD19 is a true folding innovation leading to a novel double Ig topology. We can analyze [the double Ig domain folding of CD19](https://structure.ncbi.nlm.nih.gov/icn3d/share.html?LGcQe5UM4nFx7dnL8) as formed by two Ig domains of “parallel topology”, where sequential protodomains AB-CC’ (p1/p3) and DE-FG (p2/p4) are linked by short linkers (p1-p2 and p3-p4), the two parallel domains assemble pseudosymmetrically with an inverted topology (using a membrane protein denomination) [see Figure 4]. Amazingly, in doing so it also forms two composite Ig domains, with protodomains p1 and p4 related by a local C2 symmetry, and similarly for p2 and p3. It represents a marvel of topological engineering and folding. It is worth noting that from a sequence standpoint the Ig domain formed of p2+p3 has an **inverse Ig topology** swapping sheets BED (p3) and GFCC’ (p2), as p2 precedes p3 in sequence. In essence p2 and p3 are swapped and form an inverse Ig domain [see Figure 5]. This is equivalent to a circular permutation of protodomains, yet obtained purely from folding. Ultimately, the CD19 double Ig domain forms two fused long sheets of: (BED)1(DEB)2 vs. (A’|GFCC’)1(C’CFG|A)2 [Figure 4D]. The A’|GFCC’ sheets are fused through their C’ strands, and their opposite BED sheets are fused through their D strands. A canonical IgV dimer or a swapped dimer would have GFCC’ sheets, facing each other with non bonded interactions rather than through lateral hydrogen bonded beta strands [See Figure 4F,G and Figure 6]. This is truly a remarkable structural and topological innovation.

**Figure 5.**
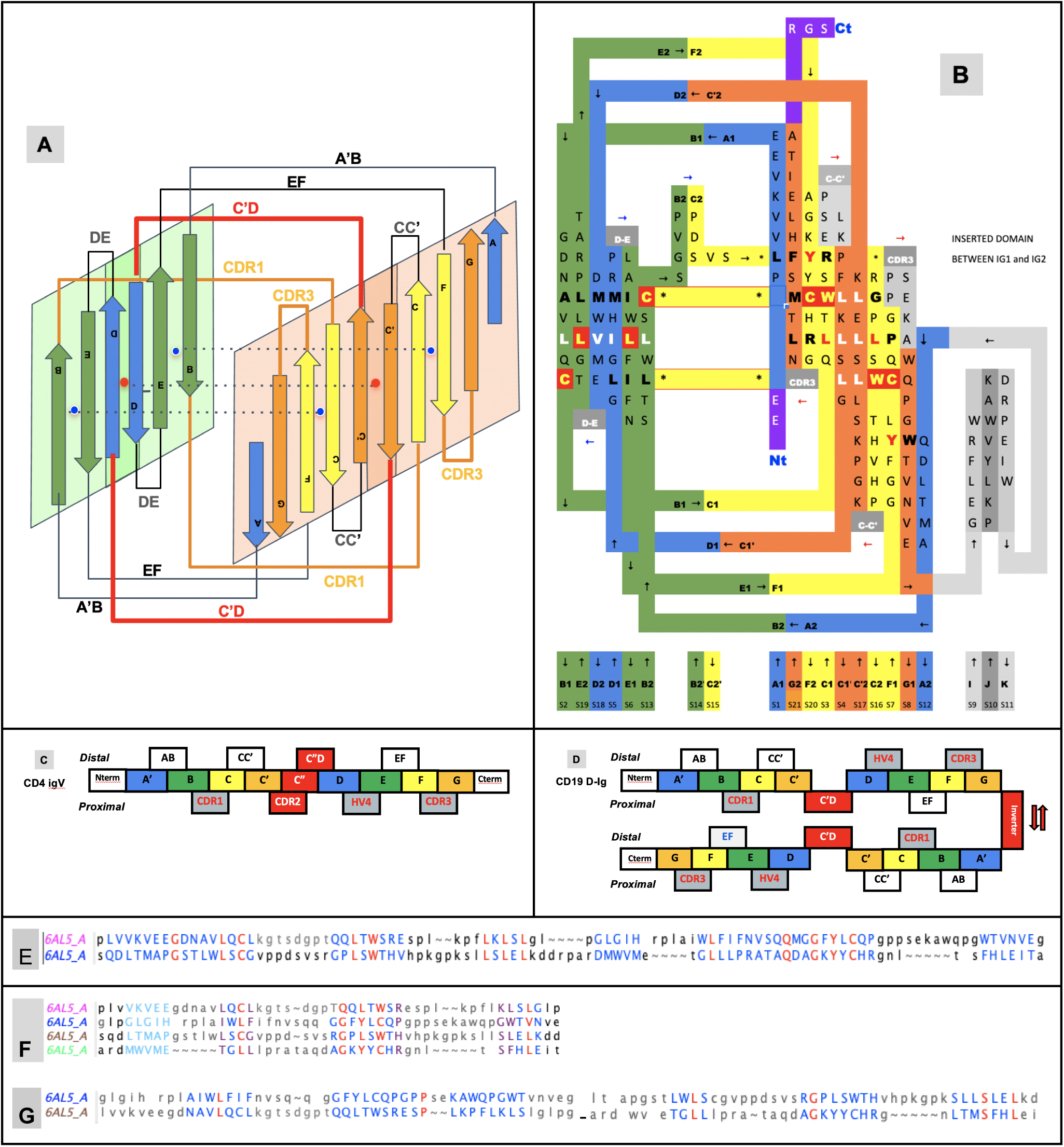
Double IgV domain topology (CD19) **A) CD19 double Ig domain topology:** Topology diagram with the two sheets facing each other. The central axis of symmetry (red dot) and the two local ones (blue dots) are shown in dashed lines. **B)** The Topology/Sequence map showing the pairing of individual residues in complementary beta strands. The comparison with the Topology/Sequence map for a single Ig domain in Figure 1D shows a conserved organization of composite Ig domains within the double domain, in particular the CCWL pattern (highlighted in red), as it is in a swapped dimer in Figure 4C. **C)** Sequence of an IgV with an A’ and no A strand as in CD4, marking the sequence stretches for strands and loops. The [CDR2 - C”- C”D loop] linker inverts protodomain 2 A’GFCC’ vs protodomin 1 BED. The CDR loops CDR1, CDR2 and CDR3 and HV4 are all on the same side, here defined as proximal for comparison. **D)** The CD19 sequence composed of protodomain 1 AB-CC’ and protodomain 2 DE-FG in parallel with a C’D short linker. In the first sequential parallel Ig domain CDR1 on the proximal side vs. CDR3 and HV4 on the distal side. The second duplicated parallel Ig domain composed of protodomain 3 and 4 is inverted through a small inserted domain (in grey in B)). This positions the two parallel Igs to intercalate and form a double Ig domain formed by two long sheets: (BED)1(DEB)2 vs. (A’|GFCC’)1(C’CFG|A’)2. In doing so two fused composite Ig domains are formed combining p1+p4 as an Ig domain of topology (AB-CC’|DE-FG) and p2+p3 as an inverted Ig domain with the topology (DE-FG|AB-CC’) since in sequence p2 (DE-FG) precedes p3 (AB-CC). The variable regions (CDR1, CDR3 and HV4) for the first composite Ig domain (p1+p4) are on the proximal side, as for a regular IgV domain in C), while in the second inverted Ig domain (p3+p4) variable regions are on the distal side. **E) Structural alignment of the two parallel Ig domains within CD19**, **i.e.** p1+p2 vs. p3+p4 matching within 1.65 A RMSD. **F) Alignment of the 4 individual Ig protodomains:** RMSD p1 vs. p2: 1.99 A, p3: 1.29 A, p4: 1.60 A on the 3 last strands BCC’ vs. EFG (since A’ strands are not structurally matched to D strands - see symmetry breaking discussion. they are indicated in light blue). **F) Alignment of the two structural Ig domains within the double Ig (CD19).** The first p1+p3 matches the second inverted Ig p2+p4 within 2.56A RMSD.

**Figure 6.**
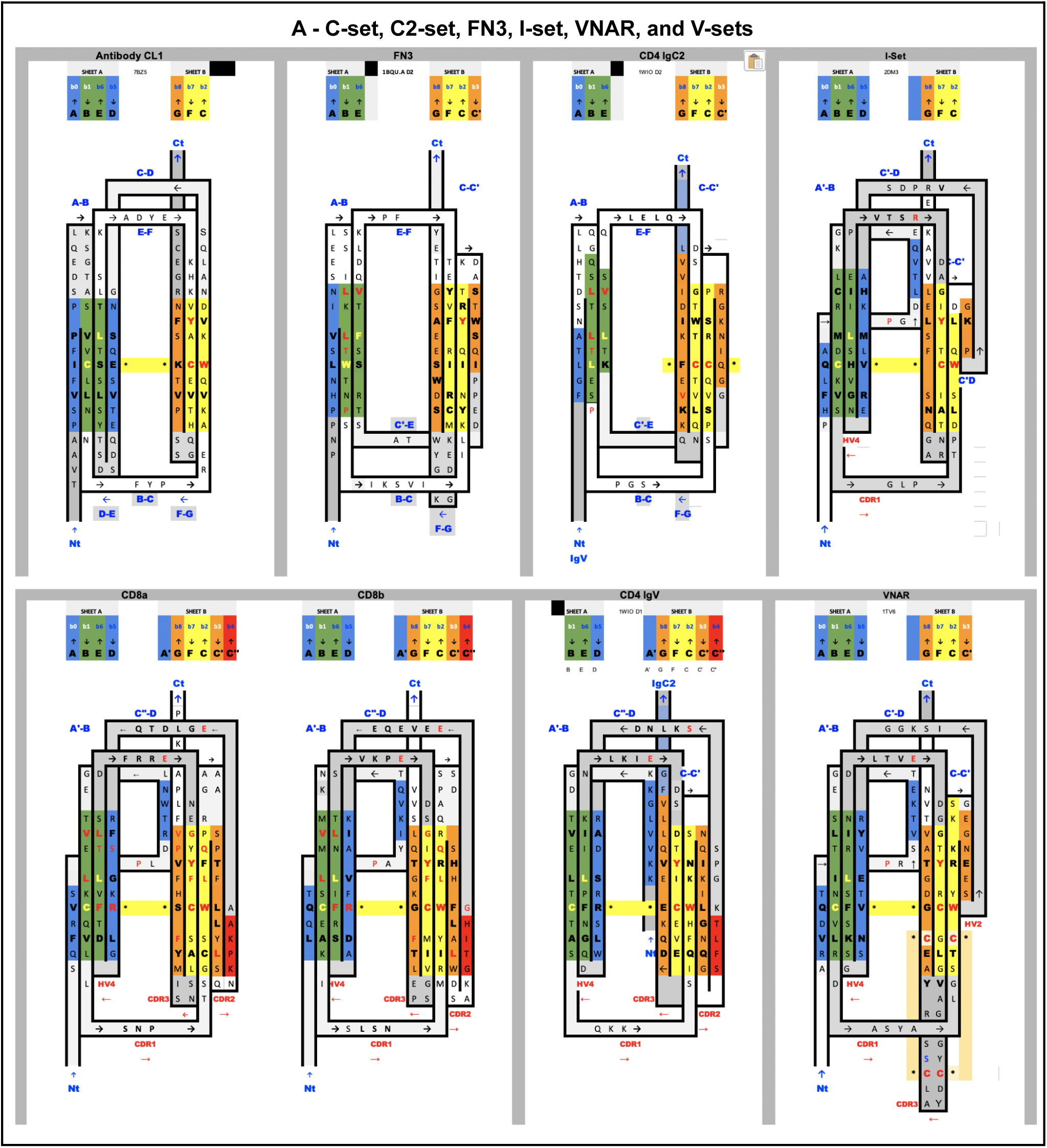
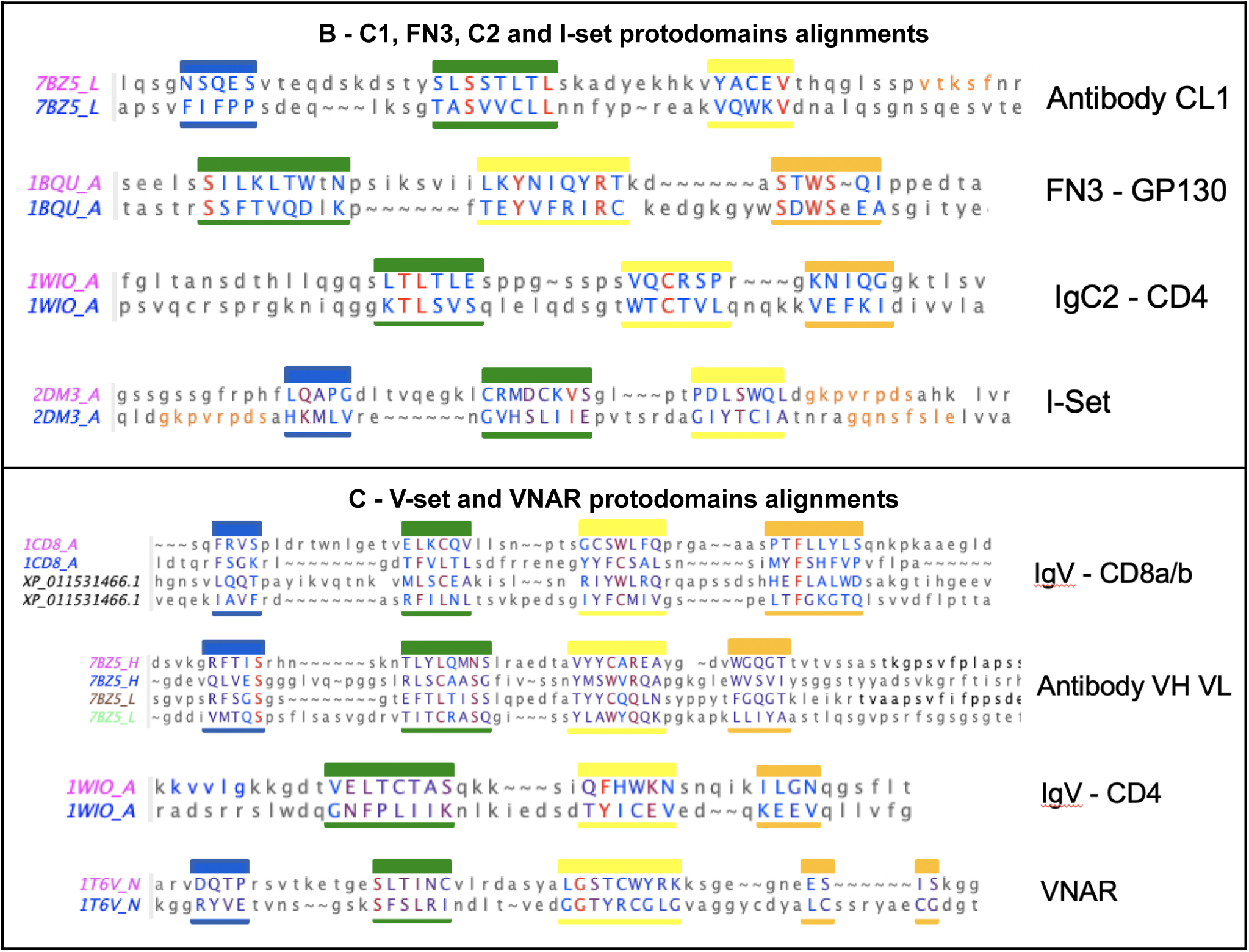
Sequence/Topology maps comparison of topological variants of Single Ig domains. **A) Sequence/Topology maps: C1-set, C2-set, FN3, I-set, VNAR** and **V-set** for CD8a/CD8b and for CD4. See text for details on differences. Capital letters are used for structurally aligned residues. Colors are the same for matching strands in protodomains: blue for strands A and D, green for B and E (a full sheet ABED will read blue-green-green-blue); yellow for strands C and F, orange for strand C’ and G, and red for C” “IgV “linkers”. Colors green and yellow of central strands highlight the self assembly of strand B-E and C-F of protodomains. FN3 and IgC2 have a similar topology, but FN3 does not share the CCW(L) present in all the selected Ig domains highlighted in central strands. Some IgV domains such as CD2 however can also miss that pattern (not shown). IgVs have a C” strand highlighted in red. We have highlighted CD8a and CD8b that form either homo or heterodimers, with a split A/A’ strand where A’ swaps from one sheet i.e antiparallel to strand B and parallel to strand G, while CD4 IgV represents another type where the strand A is only on one sheet in parallel to strand G. VNAR shares the split strand A/A’ of IgVs, which is also present in the Ig-I set. The I-set is very similar to VNARs that have only a few more residues in their HV2 region that encompass a short C’ strand antiparallel to C, and a C’-D linker that is structurally superimposable in VNAR and I-set. In fact [the overall structural alignment is very good with an RMSD of 1.83A and a surprisingly good sequence match](https://structure.ncbi.nlm.nih.gov/icn3d/share.html?c8F4iowua2TAD6ej8). **B) Alignment of protodomains in C1-, C2- and I-set domains.** Colors as in A show the corresponding strands aligning 3 out of 4 strands due to symmetry breaking (see text). In the I-set, the G strand in shown in orange as well as the would be C’ when comparing with IgVs, if it were not a protodomain “linker” in the I-set context. RMSD for protodomain alignments are for C1: 1.19 A, FN3: 1.8 A, C2: 1.98 A, I: 2.46 A (iCn3d self alignment of 2DM3) **B) Alignment of protodomains in IgVs and VNAR domains.** The symmetry breaking (see text) also extends to CD4-like IgVs where strand A is swapped in protodomain 1 vs. strand D in protodomain 2. In other IgVs such as CD8 or antibodies VH-VL domains that we added for completeness, or in VNAR the strand A is split A/A’, as in the I-set and keeps a symmetric correspondence with strand D, overall in all all 4 strands AB-CC’ vs. DE-FG, with only a partial symmetry breaking through strand A’. Protodomains alignments show consistently a structural match for conserved and co-evolved residues C in strand B and strand F vs. residues W and (most often) L in strand C and strand E respectively. RMSD for alignments are: for CD8a-IgVs 1.61 A (CD8b sequences are indicated sequence by homology with mouse CD8b structure (2ATP) whose protofdomains superimpose with and RMSD of 0.895 A and 1.54 A), followed by VH-VL protodomains for reference with an RMSD of 1.8, 0.882, 1.74 A respectively; CD4-IgV protodomains: 1.84 A; VNAR protodomains: 2.31 A

## DISCUSSION

### Structural and topological plasticity of Ig domains

The accepted nomenclature on beta strands (Lesk and Chothia 1982) has progressively been used to develop a classification of Ig-domains distinguishing four main classes of Ig domain topologies: V-set, C1-set, C2-set, I-set. All Ig domains in these classes share only four central beta-strands B,C,E,F, forming two loops straddling the two sheets of the barrel BC (CDR1) and EF, and a fifth G strand at the C terminus, forming the FG loop, called Complementary Determining Regions 3 (CDR3) in antibodies (we use the CDR nomenclature of antibodies here for all IgV domains). Two to five lateral strands A/A’ at the N terminus and C’, C’’, D that we will describe below define and distinguish these four classes (Lesk and Chothia 1982; Bork, Holm, and Sander 1994). Today many topologies and associated families have been classified and organized in a hierarchical tree (https://www.ncbi.nlm.nih.gov/Structure/cdd/cddsrv.cgi?uid=cl11960) (Lu et al. 2020). Some authors have proposed additional sets or subtypes (C3, C4, V, H, and FN3) (Halaby, Poupon, and Mornon 1999) but these complexify the topological landscape, and the more diverse the classification becomes to account for small details, the less we can see the commonalities and a possible common evolutionary scheme that led to today’s diversity in topologies. The number of all possible topological variants accounting for specific details of Ig domains in each and every functional family is quite high when considering that lateral strands can be topologically present or not, and can split and swap between the two sheets of the Ig-sandwich barrel, giving rise to a large combinatorial structural ensemble. Adding sequence diversity, one can understand the evolutionary success and diversification of these domains. Adding multiple Ig-domain chaining (tandemization) and chain oligomerization adds even more complexity and diversity.

In this paper we take a step back and look at the common symmetric architecture of all Ig domains: a protodomain decomposition allows us to reconcile all Ig-domain topologies in light of that common pseudo-symmetric domain architecture, that could imply either a variable fusion mechanism linking these protodomains together to produce various domain topologies (parallel evolution) or a divergent evolution from symmetrically fused protodomains. CD19 escapes the current classification, and exemplifies a new topological double Ig-domain innovation that can be explained by how Ig protodomains fold and combine. At the same time the protodomain decomposition validates the single Ig pseudo symmetric domain organization.

#### Ig domain classes and their topologies

All Ig domains can be related, however protodomains within Ig domains of different topologies exhibit symmetry breaking when considering AB-CC’ and DE-FG protodomains as some of the lateral strands A/A’, C’ or D are not present. Let us first go over these topological variations:

- **In IgVs**, the A strand splits in A/A’ (usually through a Proline or a set of Glycines) with **A** extending the **A|BED** sheet in antiparallel while **A’** snaps to extend the **A’|GFCC’|C’’ sheet** in parallel and straddles one lateral side on the sandwich **. In** some other IgVs such as in CD4, the A strand is reduced to A’ to participate in that sheet only. The IgV is the only Ig domain with a C’’ strand that extends the A’|GFCC’|C” sheet and forms in doing so a CDR2 loop (C’-C”). However the C” strand can also participate instead to the ABED|C”. This is the case of CTLA-4 (J. C. Schwartz et al. 2001) that has a split A/A’ on one side of the sandwich and a C” on the other side, and therefore closes the two open sides of the Ig sandwich to resemble a more a fully connected (H-bonded) 10-stranded quasi closed barrel ABEDC”C’CFGA’. In doing so, it also blocks possible lateral strand-strand (backbone level) quaternary interactions, while offering overall more surface for non-bonded quaternary interactions all around the barrel.
- In **IgC1s**, the A strand lies entirely on the **A|BED** sheet, there is no split A/A’ with A’ snapping to the other sheet which is reduced to **GFC (no C’)**
- In **IgC2s**, the A strand lies entirely on the **A|BE (no D)** sheet. There is no split A/A’ and a ***GFCC’ sheet.***
- In the **I-set,** as in many IgVs A and A’ are split and A participates to **A|BED** sheet in antiparallel while **A’** swaps to the opposing sheet in parallel to strand to G: **A’|GFCC’.** The C’ strand is very short (2-3 residues) followed by a very straight C’D linker to the D strand on the opposite sheet.
- In **VNAR** domains, the heavy chain-only variable domain of sharks (Feige et al. 2014; Greenberg et al. 1995), the topology seems to be lying between the I-set and the V-set if one considers the similarity between the protodomain linkers, i.e. with the so-called HV2 region instead of the CDR2/C” region in IgVs, that can be considered composed of a very short C’ before the C’D linker straddling to the D strand. The topological organization is similar to the I-set (Figure 6) and the C’D linker that even aligns structurally to C’D usually considered part of the HV2 variable region in e VNARs. It is tempting to see an evolutionary relationship.
- In **Cadherins** the A strand lies entirely on **A’|GFC** sheet **(no C’)** and forms a **BED** sheet.
- In **FN3**, similarly the A strand lies entirely on the **A|BE sheet (no D)** vs. the **GFC|C’** sheet.
- **In CD19** for each of the two fused structural Ig domains, the A’ strand lies entirely on **A’|GFCC’** sheets, and **BED** sheets are formed opposite to them. Both Ig-domains extend each other’s sheets in a unique way, as two fused long sheets: (A’|GFCC’)1(C’CFG|A)2 vs. (BED)1(DEB)2 that is only possible through an inversion of one domain through a unique folding/swapping mechanism [Figure 4D].

There are many papers reviewing differences between topological variants of the Ig domain (Bodelón, Palomino, and Fernández 2013; Smith and Xue 1997) and a few hypotheses have been proposed on a divergent evolution scenario between them, with the IgC2 as the oldest one (Smith and Xue 1997). The I-set (Harpaz and Chothia 1994; J.-H. Wang 2013) is intriguing as it is sometimes described as a V-set truncated on one side, missing the C’’ strand. The I-set is topologically and structurally highly similar to the VNAR domain including on strands A/A’B-CC’ and DE-FG, with a very short C’ and more surprisingly with a matching C’-D linker, missing the CDR2 loop of IgVs. VNAR however is truly a variable domain with V(D)J recombination (Greenberg et al. 1995; Ohta et al. 2019). **The four main classes of Ig domains, as well as VNAR and the FN3 domains are shown in Figure 6 as Topology/Sequence maps that visualize topological/sequence alignments, based on structure.**

Multiple cell surface receptors have an IgV-like domain at the N-terminus for ligand-binding, and many of their ligands also have an IgV-like domain at their N terminus. Ig-C1 or Ig-C2 domains often follow N-termini IgV-like domains in tandem. This is the case of CD4 (Wu et al., 1997) that shows a tandem fused IgV-IgC2 (the G strand of the IgV and the A strand of the IgC2 are fused to form a rather rigid two domains tertiary structure, that itself tandem being duplicated as 4 extracellular domains (IgV-IgC2)2. CD2 (Jones et al., 1992) also presents a tandem IgV-IgC2 yet a hinge separates the IgV G strand from the IgC2 A strand, giving the variable domain more flexibility for dimerization of the IgV domain (see swapped domain in Figure 4) and ligand interactions. All these topological variants produce sheets and side strands exposures that can lead to a variety of interaction and Ig-Ig dimer interfaces. It is beyond the scope of the present paper to review all possible interfaces, there are so many, except as it relates to canonical and inverted IgV domains and CD19 double Ig domain (see Figure 4).

#### Protodomain symmetry breaking elements

##### The protodomain linker region varies among Ig-domains

This whole **C**[c’c”d]**E** region distinguishes the various Ig-domains. They can all be reconciled by understanding the linking of two protodomains: AB-C[C’–linker–D]E-FG, forming (or removing) strands, partially breaking the symmetry and leading to topological variants. For example, [comparing the two protodomains in the I-set through a self alignment](https://structure.ncbi.nlm.nih.gov/icn3d/share.html?tBzknU8spazpwCup7), the C’ strand that would match the G strand by symmetry in an IgV (shown in orange in Figure 6) morphs into a straddling linker between the two strands sheets, between strands C and D. In FN3 domains (Campbell and Spitzfaden 1994; Youkharibache 2019) the D strand is missing, leading to a symmetric match only for strands B<>E, C<>F, C’<>G with a C’E linker [Figure 6]. Similarly, In the structurally related cadherin domains ((Shapiro et al. 1995) the C’ strand is missing, leading to a symmetric match of A<D>, B<>E, C<>F, with a CD linker [Supplement Figure]. The Ig-domain topological variants can be compared through 2D topology-sequence maps and structure based sequence alignments in Figure 6. The fusion of two protogenes and evolution would determine the length and secondary structure of protodomains linkers, and ultimately the function of the resulting Ig domain.

We cannot objectively reconstitute an old evolutionary process that gave rise to Ig domains but if we analyze multiple pseudo symmetric folds, we can infer that a substructure can duplicate with variable linker regions, these regions can be short or long and enable substructures to self assemble pseudo-symmetrically, breaking symmetry in a variety of ways. Naturally a divergent evolution scenario after duplication-fusion through insertions and deletions would also result in a variety of topologies. The two evolutive scenarios are certainly not exclusive and, in fact, likely when considering sequence similarities such as the CCW(L) signature in Ig-domains (Youkharibache 2019). In the case of a simple duplication event leading to a C2 pseudo symmetric structure, as in membrane proteins they can assemble in antiparallel to lead to a so-called inverted topology, or in parallel as in the case of CD19. Depending on the number of secondary elements in a duplicated substructure, a short or long linker may be needed to achieve a parallel or an inverted topology. A single Ig domain, when considering protodomains made of 4 strands (2 hairpins), would need a rather long linker to achieve an inverted topology of two consecutive protodomains. This is the case of an IgV domain where that linker can form a secondary structure, with a CDR2 loop between a C’ and a C” strand that complement the sheet and then a turn C”D linking across the domain to the ABED sheet. If however the linker is short or non-existent what are the choices to achieve an antiparallel protodomains organization? sacrifice either the C’ or D strand, using the sequence stretch to link the 2 sheets obtaining an IgC or an I-set (no C’) or alternatively an IgC2 or an FN3 domain. On the other hand in the case of CD19, a short linker between the A’B-CC’ (p1) and DE-FG (p2) protodomains enables a parallel organization, which can then assemble in antiparallel with a duplicated copy (p3+p4),thanks to a small inverter domain linker, placed between the two “domains” (see Figure 4).

##### The common core

The **immutable architectural element**, the common core of Ig and Ig-like domains, is the **BC-EF** intertwined symmetric straddle [Figure 1] which is at the heart of not only Ig-fold SCOP b.1, but also other folds such as SCOP folds b.2 (p53-like transcription factors) or as b.3 (prealbumin-like) (Lo Conte et al. 2000), that are all classified together as Immunoglobulin-like in other classifications (ECOD, CATH) (Dawson et al. 2017; Cheng et al. 2014). This symmetric straddle that encompasses BC (CDR1) and EF loops in symmetrically related positions is a signature of the Ig-like “inverted” topology where strands B-C and E-F substructures are intertwined in antiparallel.

##### The N terminus region: strand(s) A-A’ can also vary resulting in a partial symmetry breaking

In Ig-domains the A strand is quite interesting and also distinguishes various Ig domains. The A strand in an IgC1 will form a regular hairpin and strands AB of protodomain 1 will pair with DE of protodomain 2, but in the context of an IgV the A strand will split in two A and A’, usually through a Proline or Glycines, and the A’ will participate into the opposite sheet in parallel. In some cases the A strand lies only on the opposing sheet as A’, and the protodomains will remain pseudo symmetric only on 3 strands BED vs. CFG.

In summary, when considering Ig domains and comparing the various topologies, independently of the CDRs in antibodies, BCRs and TCRs, the most variable regions *structurally and topologically* are on the edges of the domain (barrel sandwich), on **how protodomains are linked, leading to multiple topologies** between C and E strands, with in particular the C’-CDR2-C”-C’’D structured linker in IgVs, as well as the A strand topologies variants at the N terminus: strands A (ABED sheet), or A’ or or split A-A’(A’GFCC’ sheet), participating to both sheets [see Figure 6]. Our approach sees all topological variants of the Ig domain through the lens of the **pseudo symmetric assembly of protodomains,** tied together through a **linker of variable length and secondary structure**. With the knowledge of the double Ig-fold, we have been able to rationalize protodomain assembly with that same lens, yet with a higher complexity, and we have highlighted two forms of a sequential Ig domains: with consecutive protodomains folding in parallel vs antiparallel (inverted), depending on the inter protodomain linker’s length. CD19 folding is the only instance (so far) of a double Ig fold in the structural database. This fold is unique in more than one way: in folding as parallel Ig domains, and in the pseudo symmetric assembly of these parallel Ig domains, resulting composite structural Ig domains, with an instance of an **inverse Ig-fold**, resulting from a complex protodomain swap. It is important to note that it is not a simple exchange of secondary structure elements between domains, referred to usually as “domain swap” [(Teplyakov et al. 2018; Yanshun Liu and Eisenberg 2002; Bonjack-Shterengartz and Avnir 2017)]. The Ig domain composed of structurally swapped protodomains p2+p3 is an **inverse Ig-domain,** equivalent to a circular permutation in sequence.

#### Searching for CD19 homologs in sequence or structure and for inverse Ig-domains

We performed a sequence search for a hypothetical inverse Ig domain based on a composite permuted sequence of protodomains p2-p3, but did not identify any inverse sequence, except in CD19 orthologs. If we search structurally (Holm and Laakso 2016), we get over 7000 hits however, due to the domain pseudo symmetry that matches a regular Ig domains with permuted protodomains but with a very low sequence homology, and with only 10 hits with a sequence match of above 20% identity. If we examine matches above that threshold, they also appear to be Ig domains with regular topology. The very top hit, exhibiting 23% of sequence homology, is a structural Ig domain of the Togavirin double Ig domain in the matrix remodeling associated protein 8 MRXA8 (PDBid 6JO8 and homologs). It presents what looks like a circular permutation of only half protodomain consisting of the AB strands. However, It is not a permutation at the gene level, but a domain swap of the AB substructure. In sequence, we have two chained Ig domains through an extra strand H as a linker {AB}1|{CC’C”-DE-FG)}1|**H**|{AB}2|{CC’C”-DE-FG)}2; in structure however [the two Ig structural domains](https://structure.ncbi.nlm.nih.gov/icn3d/share.html?BbigXEPw5RDi85r96) related by a C2 pseudo symmetry have the swapped topology: {CC’C”-DE-FG)}1|**H**|{AB}2 - {AB}1|{CC’C”-DE-FG)}2 as shown in Figure 4, in a head to head swapped “dimer” yet a tertiary structure (Basore et al. 2019). The first structural Ig domains appears as if it were a circular permutant in sequence with strands CC’-C”-DE-FG-**H**-AB, composed of strands **CC’-C”-DE-FG** of the first Ig sequential domain and **AB** of the second linked through a domain “linker” forming a new substructure with a new strand H between the first G strand and the second A strand in sequence, [i.e. a GH loop/strand H/HA loop linker]. The two composite structural domains use what would be the BC (CDR1) loop as a linker to swap.

##### A final note on interdigitated double domains vs. tandem domains

While unique among Ig domain topologies, the interdigitated folding of Ig protodomains in CD19 double Ig domain rather in tandem (see Figure 4) is reminiscent of that of the double Tudors. Two sequential tudor domains can swap their protodomains p1+p4 and p2+p3 *(iCn3D - 2GF7)* while tandem Tudors fold as sequentially p1+p2 and p3+p4 *(iCn3D - 1XNI) (Ying Huang et al. 2006; Youkharibache et al. 2019)* [see Supplement Figure]. This double tudor folding and the resulting topology observed in the Jumonji domain containing 2A (JMJD2A) and the Retinoblastoma-binding protein 1 (RBBP1) proteins has intrigued scientists who solved their structures (Ying Huang et al. 2006; Gong et al. 2014), yet small barrels of SH3 topology such as Tudors also have a pseudo symmetric tertiary architecture (Youkharibache et al. 2019), and their protodomains can interdigitate between consecutive sequential domains in forming pseudo symmetric double domains. As the authors conclude, “*It will be extremely interesting to understand the principle underlying the distinct folding of the double tudor domains of JMJD2A and 53BP1 despite their sequence similarity”*. In the case of small barrels with a hairpin-strand protodomain as in Tudors, it will form either tandem domains (iCn3D - tandem Tudors) or a double domain (iCn3D - double Tudor), yet in that case long links are not a geometric requirement to invert protodomains, so interdigitation of protodomains may be driven more by their sequence affinity.

In the case of CD19 however, we have proposed the hypothesis of protodomains as folding units and inter-protodomain linkers length as a distinctive element allowing Ig domains to fold in parallel (short linker) vs in antiparallel (long linker); in the latter case enabling structural tandem domain formation, while in the former enabling interdigitation. If protodomains form stable supersecondary structures, then the linker length will enable either structural tandem or interdigitated domain formation. Pseudo-symmetric assembly of protodomains as a domain requires a linker whose length depends on their folded topology. In the case of Igs with 2-hairpin protodomains, a long enough linker is needed to invert the second protodomain vs. the first one as a single Ig, but a very short linker will prohibit two consecutive protodomains from folding as a closed Ig-domain and lead to an open parallel Ig domain that can interdigitate with a copy of itself, as in CD19. Linker length is known to control intrachain domain pairing of VH and VL domains as either single chain Fv fragments vs. interchain dimeric assembly as diabodies (Poljak 1994; Holliger, Prospero, and Winter 1993). The same principle of pseudosymmetric assembly of domains is observed in the pseudosymmetric protodomain assembly, forming either tandem domains or interdigitated domains.

## METHODS

Methods have been published previously (Youkharibache 2019). Software Programs used in this work are iCn3D (J. Wang et al. 2020), CEsymm (Myers-Turnbull et al. 2014), SYMD (Tai et al. 2014) that has been integrated recently to iCn3D as a web service, Cn3D for interactive multiple structure alignment (Y. Wang et al. 2000) as well as NCBI structure databases and structure comparison servers (MMDB/VAST/VAST+) (Madej et al. 2014), the PDB servers (Burley et al. 2017), DALI server for structure searches (Holm and Laakso 2016), BLAST for sequence searches (Altschul et al. 1990), UNIPROT (UniProt Consortium 2018), and NCBI CDD annotations (Lu et al. 2020) for structural and evolutionary related annotations, in particular the latest [IgV-set CDD](https://www.ncbi.nlm.nih.gov/Structure/cdd/cddsrv.cgi?hslf=1&uid=409355), [IgC1-set CDD](https://www.ncbi.nlm.nih.gov/Structure/cdd/cddsrv.cgi?uid=409354) and many instances of [IgC2 CDD](https://www.ncbi.nlm.nih.gov/Structure/cdd/cddsrv.cgi?hslf=1&uid=409491) and [IgI CDD](https://www.ncbi.nlm.nih.gov/Structure/cdd/cddsrv.cgi?hslf=1&uid=409545) with detailed Ig sequence-structure features, down to each and every strand of these domains.

## CONCLUSION

The analysis of pseudosymmetric domains, while not demonstrating a folding pathway, demonstrates stable folded elements that we call protodomains. The Ig domain can be considered formed of two protodomains and the recent identification of a double Ig domain present in CD19 gives a confirmation of the protodomain hypothesis and shows the ability of Ig-protodomains to fold independently and assemble in either parallel or antiparallel, similarly to membrane proteins of parallel vs inverted topologies. We have also seen the propensity of Ig domains to pair symmetrically in either parallel or inverted “quaternary topologies’’ as a result of the tertiary pseudo symmetry of Ig-domains themselves.

The evolutionary origin of the Ig domains of varying topology is not entirely solved by this decomposition, yet the decomposition in protodomains involving a symmetric combination as well as symmetry breaking elements offers a number of possible scenarios throughout evolution to explain these topological Ig-domain variants. This also tends to suggest that the IgV may be the ancestral topological form of single Ig-domains, but the double Ig-domain would have had to have its own path.

Importantly, the pseudo-symmetric deconstruction of Ig-domains provides a frame of reference for innovative molecular design. We have seen the formation of interdigitated protodomains in double domains, the largely unexplored possibilities of linkers to control folding. We have observed an inverse Ig domain within the double Ig domain, and one can extrapolate to possibly engineer (circularly) permuted Ig domains for targeted application. As nanobodies applications expand, harnessing the pseudosymmetry of single Ig domains can offer multivalent binding surfaces, for cell surface ligand designs efficient targeting of Ig-based cell surface receptors, to lead to next generation checkpoint inhibitors and chimeric antigen receptors for CAR T-cell therapies in the burgeoning field of immunoengineering.

## ACKNOWLEDGEMENTS

Special thanks to Jiyao Wang (NCBI) for a wonderful collaboration in the development of iCn3D that illustrate this paper all along with 3D visualization through lifelong hyperlinks; James Song (NCBI) for the new CDD structural annotations of Ig domains and his feedback, Tom Madej (NCI/NCBI) for his development and support of VAST and VAST+ structural database, used thoroughly in structural analyses; Aleix Lafita (EBI) and Spencer Bliven (Paul Scherrer Institut) for their development and support of CE-symm used to identify tertiary and quaternary symmetries in proteins; Emily Tai (NCI) and BK Lee (NCI) for making SYMD available to set up a web service to annotate symmetry in iCn3D; Raul Cachau (FNLCR) for sharing his very deep knowledge and insight on protein structure and structural analysis, Peter Rose (SDSC) and Jose Duarte (RCSB) for their help in navigating protein symmetry annotations in the PDB.

This work has been funded by the NIH intramural program.

